# Inversions shape the divergence of *Drosophila pseudoobscura* and *D. persimilis* on multiple timescales

**DOI:** 10.1101/842047

**Authors:** Katharine L Korunes, Carlos A Machado, Mohamed AF Noor

**Affiliations:** Department of Evolutionary Anthropology, Duke University, Durham, NC 27708; Department of Biology, University of Maryland, College Park, Maryland 20742; Biology Department, Duke University, Durham, NC 27708

**Keywords:** inversions, introgression, divergence, recombination, speciation

## Abstract

By shaping meiotic recombination, chromosomal inversions can influence genetic exchange between hybridizing species. Despite the recognized importance of inversions in evolutionary processes such as divergence and speciation, teasing apart the effects of inversions over time remains challenging. For example, are their effects on sequence divergence primarily generated through creating blocks of linkage-disequilibrium pre-speciation or through preventing gene flux after speciation? We provide a comprehensive look into the influence of chromosomal inversions on gene flow throughout the evolutionary history of a classic system: *Drosophila pseudoobscura* and *D. persimilis*. We use extensive whole-genome sequence data to report patterns of introgression and divergence with respect to chromosomal arrangements. Overall, we find evidence that inversions have contributed to divergence patterns between *Drosophila pseudoobscura* and *D. persimilis* over three distinct timescales: 1) pre-speciation segregation of ancestral polymorphism, 2) post-speciation ancient gene flow, and 3) recent gene flow. We discuss these results in terms of our understanding of evolution in this classic system and provide cautions for interpreting divergence measures in similar datasets in other systems.

## Introduction

Divergence and speciation sometimes occur in the presence of gene exchange between taxa. Estimates suggest that over 10% of animal species hybridize and exchange genes with related species (Mallet 2005). Analyses in the genomic era have provided further evidence of the widespread prevalence of hybridization and revealed many previously unanticipated instances of hybridization (Payseur & Rieseberg 2016; Taylor & Larson 2019). Understanding genetic exchange between species gives us insights into the genetic processes underlying later stages of the speciation continuum. Many approaches can examine evidence for introgression, including comparing sympatric vs. allopatric populations to test for differences in nucleotide divergence. Other available methods for characterizing gene flow include model-based frameworks and examinations of differences in divergence reflected in coalescence times. Differences in coalescence times are often observed between species in regions where recombination is limited in hybrids, such as fixed chromosomal inversion differences (Guerrero *et al.* 2012). When species differing by inversions hybridize, the collinear genomic regions can freely recombine, while inverted regions experience severely limited genetic exchange in hybrids and often accumulate greater sequence differentiation over generations. This process can lead to locally adapted traits and reproductive isolating barriers mapping disproportionately to inverted regions (reviewed in Ayala & Coluzzi 2005; Butlin 2005; Jackson 2011).

Many studies examine the timing and frequency of gene exchange between hybridizing species, with emphasis on the implications of patterns of divergence in allopatric vs sympatric pairs and in regions of reduced recombination in hybrids. However, different approaches sometimes yield distinct interpretations regarding the presence or extent of introgression. Model-based approaches yield important insights but are also limited in the scenarios that they consider and the assumptions they make about population histories and evolutionary rates (reviewed in Payseur & Rieseberg 2016). Further, shared patterns of variation are often interpreted as evidence of ongoing gene flow, but segregating ancestral polymorphism could also be the primary, or even the sole, driver of these patterns (Fuller *et al.* 2018). In the ancestral population of two species, segregating chromosomal inversions may shield inverted regions of the genome from recombination, thus facilitating the divergence of sympatric ecotypes or populations. Heightened within-species differentiation in inverted regions has been observed in many systems, including *Rhagoletis pomonella* (Michel *et al.* 2010), *Anopheles gambiae* (Manoukis *et al.* 2008), and *Mimulus guttatus* (Lowry & Willis 2010). Such heightened differentiation between karyotypes may persist along the speciation continuum, making it difficult to disentangle the effects of inversions reducing recombination in the ancestral population vs reducing introgression upon secondary contact. Fuller *et al*. (2018) recently discussed the possibility that ancestrally segregating inversions that sort between species may provide a "head-start" in molecular divergence, possibly predisposing them to harbor a disproportionate fraction of alleles associated with species differences. Unlike models assuming homogenization of collinear regions via post-speciation gene flow, this model predicts that young species that diverged in allopatry may also exhibit higher divergence in inverted regions than collinear regions. These models are not mutually exclusive: dynamics of the ancestral population as well as post-speciation gene flow can shape patterns of variation between species.

Disentangling the effects of ancestral polymorphism from the effects of post-speciation gene flow is a fundamental puzzle in understanding speciation. To achieve a cohesive picture of how hybridization influences divergence and speciation, we need to consider the approaches outlined above in a model system with extensive whole-genome sequence data to assess models and reconcile interpretations of possible signals of introgression. The sister species pair *Drosophila pseudoobscura* and *D. persimilis* present an ideal opportunity to dissect an evolutionary history of divergence nuanced by multiple inversions, lineage sorting, and gene flow. Despite the rich history of work on understanding speciation and divergence in *D. pseudoobscura* and *D. persimilis*, there are unresolved questions about the rates and timing of introgression between these species. A few F_1_ hybrids of these species have been collected in the wild (Powell 1983) and many previous studies have documented molecular evidence of introgression, detectable in both nuclear and mitochondrial loci (e.g., Machado *et al.* 2002; Machado & Hey 2003; Hey & Nielsen 2004; Fuller *et al.* 2018). Inverted regions between these species exhibit greater sequence differences than collinear regions, and this pattern was previously inferred to result from introgression post-speciation. In the largest scale study, McGaugh and Noor (2012) used multiple genome sequences of both species and an outgroup, and reinforced previous studies (e.g., Noor *et al*. 2007) showing that the three chromosomal inversions differ in divergence time. They inferred a "mixed mode geographic model" (Feder *et al.* 2011) with sporadic periods of introgression during and after the times that the inversions spread. However, in addition to confirming evidence for gene flow between *D. pseudoobscura* and *D. persimilis* after speciation, Fuller *et al.* (2018) recently argued the inversions arose within a single ancestor species, differentially sorted in the descendant species, and this sorting of ancestral polymorphisms may explain observed patterns of nucleotide variation. To fully understand the role of hybridization in the speciation process, the contrasting models must be reconciled.

We acquired extensive whole-genome sequence data to re-explore patterns of introgression and divergence in the *Drosophila pseudoobscura* / *D. persimilis* system. We leverage the allopatric *D. pseudoobscura* subspecies, *D. pseudoobscura bogotana* (*D. p. bogotana)* and two outgroup species (*D. miranda* and *D. lowei*) to distinguish recent from ancient effects of inversions on gene flow. To help clarify the role of inversions over the evolutionary history of these species, we consider three distinct time scales: 1) pre-speciation segregation of ancestral polymorphism, 2) post-speciation ancient gene flow, and 3) recent introgression (Figure 1). Patterns of divergence between *D. persimilis* and allopatric *D. p. bogotana* can be explained by the effects of segregating ancestral polymorphism and by gene flow prior to the split of *D. p. bogotana* (Figure 1, green and blue regions). In comparing the sympatric species, *D. persimilis* and *D. p. pseudoobscura*, the same forces factor into patterns of divergence, with the added effects of recent or ongoing gene flow (Figure 1, orange arrows). We leverage these two comparisons to weigh the relative contributions of recent genetic exchange.

**Figure 1.**
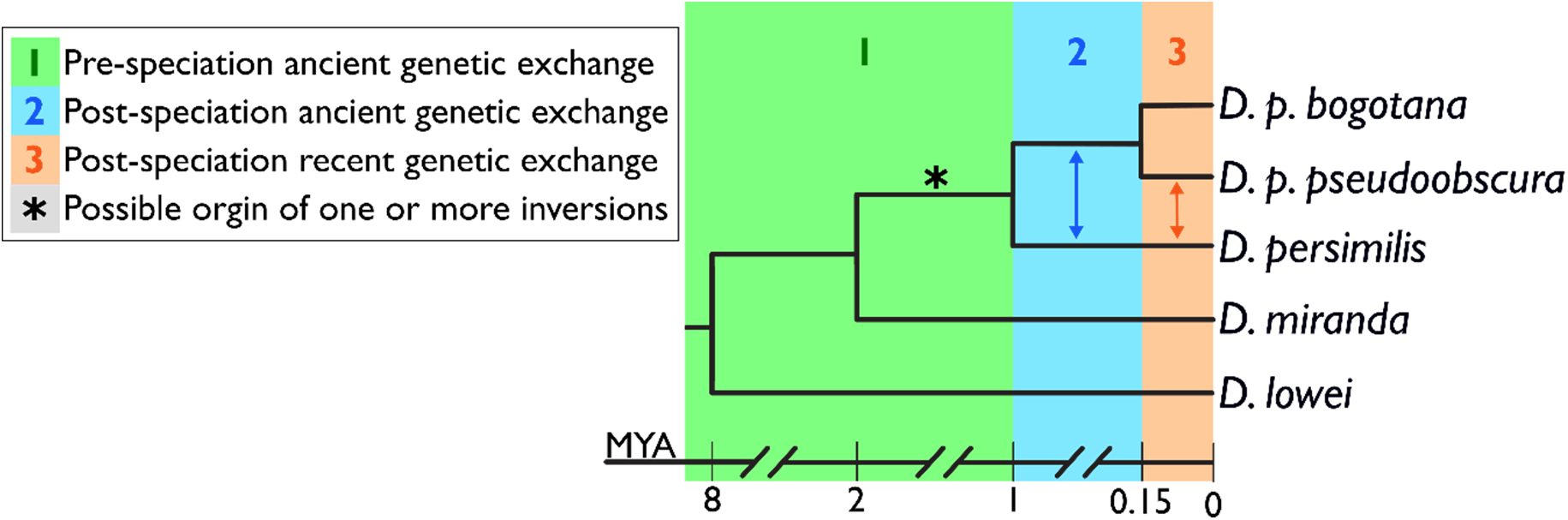
Gene flow in the context of the evolutionary history of *D. pseudoobscura* and *D. persimilis*. We consider how inversions differing between *D. pseudoobscura* and *D. persimilis* might shape patterns of divergence by affecting gene flow at 3 timescales: 1) pre-speciation recombination in ancestral populations with segregating inversion polymorphisms, 2) post-speciation ancient gene flow, and 3) recent introgression. Here, we show the evolutionary context and approximate divergence times of the taxa considered in the present study, with arrows indicating post-speciation gene flow between *D. pseudoobscura* and *D. persimilis.*

We first examine patterns of divergence in inverted regions compared to collinear regions, and we discuss evidence for early, pre-speciation exchange. We next examine evidence of post-speciation gene flow to test whether some of this genetic exchange predates the split of *D. p. bogotana*, and we discuss signals of possible introgression in the past 150,000 years since the split of the allopatric *D. p. bogotana*, from North American *D. pseudoobscura* (*D. p. pseudoobscura*) (Schaeffer & Miller 1991). We implement Patterson’s D-statistic to contrast the sympatric (*D. p. pseudoobscura*) and allopatric (*D. p. bogotana)* subspecies in their similarity to *D. persimilis*. Kulathinal *et al.* (2009) previously argued that recent post-speciation gene flow contributes to the difference in coalescence time between inverted and collinear regions, observable in the higher genetic similarity in collinear regions between *D. persimilis* and sympatric *D. p. pseudoobscura* compared to similarity between *D. persimilis* and allopatric *D. p. bogotana*. That study also tested for an excess of shared, derived bases between *D. persimilis* and *D. p. pseudoobscura* compared to *D. persimilis* and allopatric *D. p. bogotana.* Their application of this D-statistic precursor suggested a borderline statistically significant signature of gene flow, but this test was limited by low sequencing coverage. Nonetheless, consistent with that and other previous studies, we observe higher divergence between these species in inverted than collinear regions, and our implementation of Patterson’s D-statistic indicates very recent gene exchange in collinear regions. Notably, divergence measures to *D. persimilis* are also higher for allopatric than sympatric *D. pseudoobscura* subspecies in *both* inverted and collinear regions. One possible explanation for this pattern is extensive recent gene exchange throughout much of the genome, even in inverted regions. To determine the relative effect of introgression, we correct for different evolutionary rates across taxa, and we consider the role of segregating inversions in ancestral populations. We discuss these results in the context of the extensive past work towards understanding divergence and speciation in this classic system, and we provide cautions for interpreting divergence measures in similar datasets in other systems.

## Methods

### Genomic datasets

Whole-genome short-read sequence data was analyzed from 19 *D. p. pseudoobscura* and 8 *D. persimilis* strains, along with 4 *D. p. bogotana* strains as an allopatric point of comparison (both males and females were sequenced, all from inbred strains listed in Supplementary Table 1 with SRA accessions). We used *D. lowei* as an outgroup. *D. lowei* and *D. pseudoobscura* likely diverged 5-11 MYA (Beckenbach *et al.* 1993), and hybrids between these two species are sterile (Heed *et al.* 1969). Scripts used for genome alignment, SNP calling, and analyses are available on GitHub (https://github.com/kkorunes/Dpseudoobscura_Introgression). To avoid biasing identification of variants towards *D. pseudoobscura*, the *D. miranda* reference genome assembly (DroMir2.2; GenBank assembly accession GCA_000269505.2) was chosen as the reference for all subsequent alignments. *D. miranda* diverged from *D. pseudoobscura* only within the past ~2 million years (Wang & Hey 1996), facilitating alignment of *D. pseudoobscura* and *D. persimilis* genomes to the *D. miranda* genome assembly. Further, the arrangement of the assembled *D. miranda* chromosomes matches the published contig order and arrangement of *D. pseudoobscura* (Schaeffer *et al.* 2008), but with the advantage of being assembled into 6 continuous chromosome arms: chromosomes XL, XR, 2, 3, 4, and 5. Here, we analyze the majority (83%) of the assembled genome. We exclude only regions where we cannot reasonably examine introgression and divergence: chromosome 3, which presents confounding factors from its inversion polymorphisms within species (Dobzhansky & Epling 1944; Powell 1992), and the very small (<2 Mb) portion of the genome found on the largely nonrecombining “dot” chromosome (chromosome 5).

### Alignments and variant calling

To confirm the arrangement of *D. pseudoobscura* contigs with respect to the *D. miranda* reference, each *D. pseudoobscura* chromosome was split into lengths of 1 Mb, and these segments were aligned to the *D. miranda* reference using BWA-0.7.5a (Li & Durbin 2009). We then extracted the 2 kb regions surrounding published inversion breakpoints to obtain the breakpoint locations in the coordinates of the *D. miranda* reference (see Supplementary Table 2). After confirming that the arrangement of the assembled *D. miranda* chromosomes matched the arrangement of the *D. pseudoobscura* contig order and arrangement described by Schaeffer *et al.* (2008), all sequencing data were aligned to the reference genome of *D. miranda* using BWA-0.7.5a (Li & Durbin 2009), and Picard was used to mark adapters and duplicates (http://broadinstitute.github.io/picard). Variants were called used GATK v4 and filtered based GATK’s hard filtering recommendations (McKenna *et al.* 2010; Van der Auwera *et al.* 2013), excluding sites with QualByDepth (QD) < 2.0, FisherStrand (FS) > 60, StrandOddsRatio (SOR) > 3.0, MQ < 40, MQRankSum < −12.5, ReadPosRankSum < −8.

### Patterns of divergence

The resulting VCF files were then processed using PLINK (Purcell *et al.* 2007). VCFs were converted to PLINK’s bed/bim format, keeping only sites that passed the filters described above. SNPs were pruned for linkage disequilibrium using the --indep-pairwise function of PLINK (“--indep-pairwise 50 50 0.5”) before performing principal components analysis (PCA) using PLINK’s --pca function to confirm the grouping of individuals within their respective species (Figure 1; Supplementary Figure 1). Admixtools was used to implement Patterson’s D-statistic (Patterson *et al.* 2012) using *D. lowei* as on outgroup to polarize ancestral vs derived alleles. For input into Admixtools, we used the *convertf* program of Admixtools to convert each PLINK ped file to Eigenstrat format which includes a *genotype* file, a *snp* file, and an *indiv* file. Per recommendations in the Admixtools documentation, we defined the physical positions of each SNP in the *snp* file to be 10 kb apart from each adjacent SNP to allow Admixtools to interpret every 100 SNPs as 1 Mb or 1cM, since this software uses centiMorgans as the unit for block size during jackknifing and assumes that 1 Mb = 1 cM. We set the block size parameter to 0.01 cM, which in this case is interpreted as blocks of 100 SNPs. Next, *qpDstat* was used to obtain D-statistics for each chromosome. These four-population tests were of the form (((A,B), C), D), where A = *D. p. bogotana*, B = *D. p. pseudoobscura*, C = *D. persimilis*, and D = *D. lowei.* To study signatures of introgression along the genome, we applied f_d_ (Martin *et al.* 2015) in genomic intervals that presented an excess of ABBA over BABA sites. Using non-overlapping windows of 100 SNPs, we calculated f_d_ using the

Dinvestigate program from Dsuite (Malinsky *et al.* 2020). Absolute divergence, D_xy_, was calculated using custom scripts over fixed window sizes of 50 kb. D_xy_ was calculated from variant and invariant sites after subjecting SNPs to the filters described above and filtering invariant sites based on depth (depth >= 10). Per-site depths for all sites were acquired from BAM files using Samtools (“samtools depths -a <in>”) (Li *et al.* 2009). Finally, to test for differences in evolutionary rate that might influence observed patterns of divergence and gene flow, differences in substitution rates among the lineages were assessed with Tajima’s relative rate test, using *D. lowei* as the outgroup (Tajima 1993; scripts for implementation available on GitHub repository linked above). Tajima’s relative rate test was applied to the combined set of SNPs from chromosomes 2, 4, XL, and XR—excluding sites where the outgroup *D. lowei* was heterozygous or missing data.

### Models of gene flow

To test for evidence of gene flow after the split of *D. pseudoobscura* and *D. persimilis*, but before the split of *D. p. bogotana*, we used the maximum-likelihood methods derived by Costa & Wilkinson-Herbots (2017) to compare three scenarios of divergence between *D. persimilis* and *D. p. bogotana* (Figure 2): (A) divergence in isolation (Iso) without gene flow following the split of an ancestral population; (B) divergence in isolation-with-migration (IM) with constant (but potentially asymmetric) gene flow since the split of an ancestral population until the present; and (C) divergence in isolation-with-initial-migration (IIM) with gene flow until some timepoint in the past and divergence in isolation since that timepoint. Under the IIM model, we tested four scenarios: IIM1 estimates parameters under divergence with potentially asymmetric bidirectional gene flow until some timepoint in the past, assuming constant population sizes; IIM2 is the same scenario as IIM1, but allows for changes in population sizes; IIM3 and IIM4 are the same as IIM2, but assume unidirectional gene flow from population 2 to population 1 (IIM3) or from population 1 to population 2 (IIM4).

**Figure 2.**
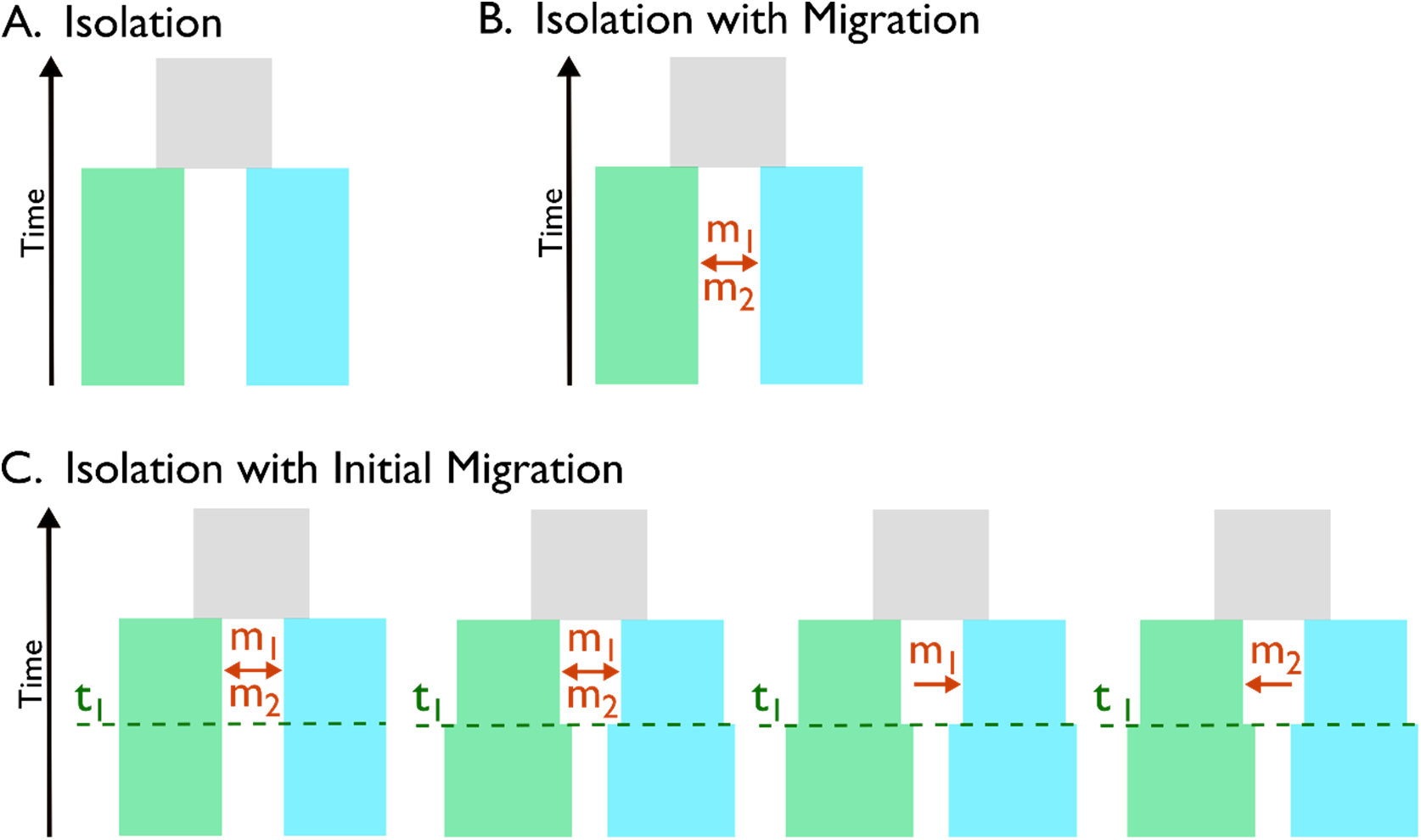
Models of Divergence. We considered the following coalescent models described by Costa & Wilkinson-Herbots (2017) to consider scenarios of divergence of *D. persimilis* and *D. p. bogotana* since the split of the ancestral population (gray box): (A) divergence in isolation without gene flow; (B) divergence in isolation-with-migration (IM) with constant (but potentially asymmetric) gene flow; and (C) divergence in isolation-with-initial-migration (IIM) with gene flow until some timepoint (t_1_) in the past. Under the IIM model, we tested the four scenarios shown from left to right: the first scenario assumes constant population sizes, the second allows for changes in population sizes, and the third and fourth allow for changes in population size but assume unidirectional gene flow.

We computed the likelihood of our *D. persimilis* and *D. p. bogotana* sequence data under each of the six scenarios described above and in Figure 2 (Iso, IM, IIM1, IIM2, IIM3, and IIM4). To reduce potential effects of selection, we used intergenic loci spaced at least 2 kb apart, similar to the strategy of Wang & Hey (2010). Linkage-disequilibrium decays within tens to hundreds of bases in *Drosophila* (Langley *et al.* 2000), so we expect that avoiding genic regions will minimize the effects of linked selection. To identify intergenic regions in the *D. miranda* genome, we used the set of all *D. pseudoobscura* gene annotations published by Flybase (http://flybase.org, Full Annotation Release 3.04), and we used BLAST to identify genomic regions with significant similarly to the *D. pseudoobscura* gene annotations, using cutoffs of evalue = 10^−6^ and percent identity = 80 (Altschul *et al.* 1990). From the remaining regions, we then randomly sampled 500 bp segments separated by at least 2 kb to create a set of ~15,000 intergenic loci. We then randomly divided these loci into three nonoverlapping subsets to satisfy the models’ requirement of independent estimates of pairwise differences and mutation rates in loci (1) within *D. persimilis*, (2) within *D. p. bogotana*, and (3) between *D. persimilis* and *D. p. bogotana.* Costa & Wilkinson-Herbots (2017) recommends using per-locus relative mutation rates, which we calculated using the average distance to the outgroup *D. lowei*, following the equation from Yang (2002), which gives the relative mutation rate at a locus as the outgroup distance at that locus divided by the average outgroup distance along all loci. To select the model that best fits the data, we then tested the relative support among the divergence models using likelihood-ratio tests following the sequence of pairwise comparisons shown in Table 1, where the degrees of freedom in each test is the difference in the dimensions of parameter space (Costa & Wilkinson-Herbots 2017). To ensure that our results were robust against the effects of linkage within inverted regions, we next sampled regions expected to be freely-recombining throughout the timescales examined: i.e., we repeated these analyses excluding any loci from the inverted regions, leaving ~11,000 intergenic, collinear loci.

**Table 1.** Forward selection of the best model^†^ of *D. persimilis - D. p. bogotana* divergence using the maximized log-likelihood (LogL) under each model in likelihood-ratio tests.

| H<sub>0</sub> | H<sub>1</sub> | Deg. of Freedom | LogL H<sub>0</sub> | LogL H<sub>1</sub> | LRT Statistic | P-value |
| --- | --- | --- | --- | --- | --- | --- |
| Iso | IM | 2 | -33702.73 | -33659.39 | 86.68 | 1.505e-19 |
| IM | IIM1 | 1 | -33659.39 | -33659.39 | 0 | - |
| IM | IIM2 | 3 | -33659.39 | -33538.92 | 240.94 | 5.960e-52 |
| IIM1 | IIM2 | 2 | -33659.39 | -33538.92 | 240.94 | 4.792e-53 |
| IIM1 | IIM3 | 1 | -33659.39 | -33543.76 | 231.26 | 3.166e-52 |
| IIM1 | IIM4 | 1 | -33659.39 | -33542.89 | 233 | 1.322e-52 |

<sup>†</sup> See Figure 3 for illustration of the different models.

## Results

### Patterns of divergence in inverted vs collinear regions

The suppression of crossing over within inversions leads to distinct signatures of nucleotide divergence within and near inversions. Figure 3 presents windowed (50 kb windows) estimates of divergence between *D. persimilis* and *D. p. bogotana* and divergence between *D. persimilis* vs *D. p. pseudoobscura* for the three chromosome arms that contain fixed (chromosome 2, XL) or nearly-fixed (chromosome XR) inversion differences between *D. persimilis* and *D. pseudoobscura*. As previously observed, divergence is low in regions near the centromere (Noor *et al.* 2007; Kulathinal *et al.* 2009). To consider the effects of inversions on divergence, we contrast observed patterns within inversions to regions outside the inversions (collinear) and to the subset of collinear regions that can be predicted to be reasonably freely-recombining (denoted as collinear_FR_). Collinear_FR_ excludes the 5 Mb windows adjacent to telomeric and centromeric ends of chromosome assemblies, which undergo very little crossing over and harbor reduced sequence diversity (Andolfatto & Wall 2003; Kulathinal *et al.* 2008; Stevison & Noor 2010). Similarly, collinear_FR_ excludes regions within 2.5 Mb outside of inversion breakpoints, based on previous studies which have reported suppression of crossing over extending 1-2 Mb beyond inversion breakpoints (Machado *et al.* 2007; Kulathinal *et al.* 2009; Stevison *et al.* 2011).

**Figure 3.**
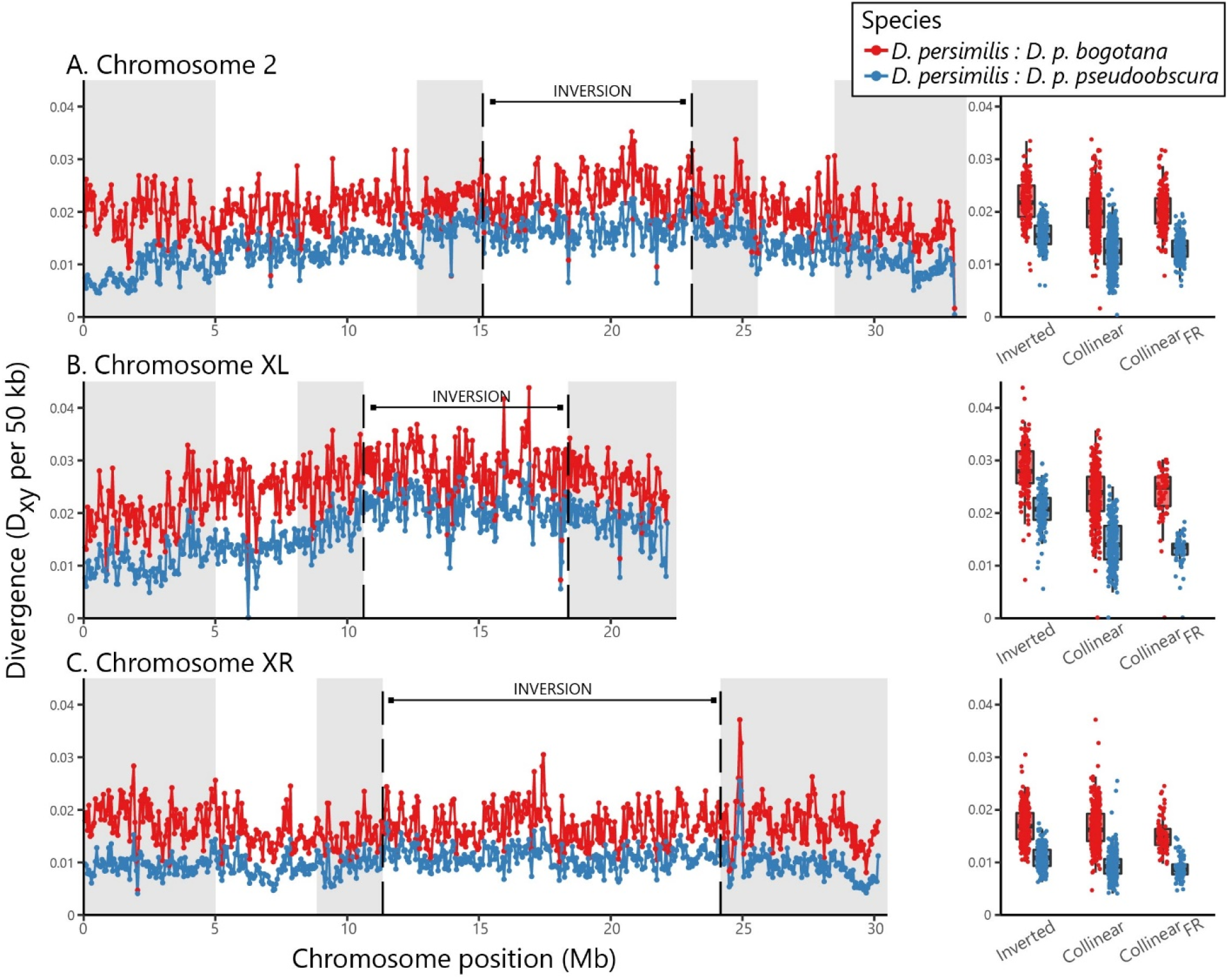
Genome-wide divergence between species. On the left, each of the 3 inversion-bearing chromosome arms are plotted from centromere (0) to telomere, inversion boundaries are shown with vertical black lines. D_xy_ per 50 kb window is plotted to show divergence between *D. persimilis* and *D. p. bogotana* (red) and divergence between *D. persimilis* vs *D. p. pseudoobscura* (blue). Boxplots (right) summarize these divergence estimates by region: Inverted, Collinear, and Collinear_FR_. Collinear_FR_ is the subset of collinear positions predicted to be freely recombining (excludes the grayed-out positions near inversion breakpoints or chromosome ends).

Confirming many previous studies (Machado *et al.* 2007; Noor *et al.* 2007; Kulathinal *et al.* 2009; Stevison *et al.* 2011; McGaugh & Noor 2012), we observed that *D. persimilis* vs *D. p. pseudoobscura* divergence is significantly higher in inverted than collinear windows, regardless of whether the inverted regions are compared to all collinear windows or to the collinear_FR_ subset (see Supplementary Table 3 for average divergence estimates and statistical comparisons). *D. persimilis* vs allopatric *D. p. bogotana* divergence is also higher in inverted than collinear windows on chromosomes 2 and XL, with a nonsignificant difference on chromosome XR (though the difference is significant if the comparison is restricted to collinear_FR_; Supplementary Table 3). The higher divergence in inverted vs collinear regions could be due to pre-speciation segregation of inversion polymorphisms in the ancestral population or to interspecies gene flow homogenizing collinear regions. We next examined evidence of pre-speciation genetic exchange.

### Confirming evidence for exchange between karyotypes early in the speciation continuum

Previous studies identified differences in sequence divergence between *D. persimilis* and *D. p. pseudoobscura* among the three inverted regions (Noor *et al.* 2007; McGaugh & Noor 2012; Fuller *et al.* 2018). They interpreted this finding as evidence that the derived inversions arose during, or interspersed by, periods where gene exchange was occurring in other regions of the genome. For both *D. persimilis* vs *D. p. pseudoobscura* and *D. persimilis* vs *D. p. bogotana*, we compared measures of windowed divergence among the inverted regions. Each pairwise comparison between the inversions yielded a significant difference wherein XL > 2 > XR (p < 0.0001, Mann-Whitney U test; Supplementary Table 3). Our observed average divergence within the inversions confirms previous accounts of the relative divergence and apparent relative age of the inversions (Noor *et al.* 2007; McGaugh & Noor 2012; Fuller *et al.* 2018), supporting the previous suggestions of ancestral exchange prior to the completion of speciation. We next explored the possibility of exchange after speciation but before the split of *D. p. pseudoobscura* and *D. p. bogotana* (region 2 in Figure 1).

### Evidence for early post-speciation exchange

Much of the previous support for post-speciation gene flow between these species has focused on comparisons of *D. persimilis* and *D. pseudoobscura* (Wang *et al.* 1997; Machado *et al.* 2002; Hey & Nielsen 2004; Kulathinal *et al.* 2009; Fuller *et al.* 2018). To distinguish between post-speciation ancient gene flow and recent introgression (Figure 1), we leveraged the allopatric *D. p. bogotana*. Some past studies suggesting evidence of introgression between *D. pseudoobscura* and *D. persimilis* also leveraged this allopatric subspecies (Machado & Hey 2003; Brown *et al.* 2004; Chang & Noor 2007; Kulathinal *et al.* 2009). We sampled intergenic loci from *D. persimilis* and *D. p. bogotana*, and we fit the observed patterns of nucleotide variation to models of divergence in isolation, isolation-with-migration (IM), and isolation-with-initial-migration (IIM) using maximum-likelihood estimation of parameters under these models (Figure 2; Costa & Wilkinson-Herbots 2017). In traditional IM models applied to infer gene flow, parameter estimates can be skewed by the underlying assumption that gene flow is constant. IIM specifically addresses this assumption by operating on the premise of an initial period of gene flow followed by isolation. Maximizing parameter likelihoods under an IIM framework is appropriate for the *D. persimilis* and *D. p. bogotana* comparison, given our knowledge that these taxa have been evolving in allopatry for the past 150,000 years (Schaeffer & Miller 1991). Indeed, nested model comparison to test the relative support among the models rejects the null hypothesis of divergence in isolation and suggest that IIM models best fit the data. These results were consistent between the full set of loci and the subset sampled from only collinear regions (Table 1 and Supplementary Table 4). All models allowing for migration gave a significantly better fit than a model of strict divergence in isolation, and the log-likelihood of the data under the tested models was maximized in the IIM2 scenario (Table 1 and Supplementary Table 4, 5). The IIM2 model estimates parameters under divergence with potentially asymmetric bidirectional gene flow until some timepoint in the past and, unlike the IIM1 model, does not assume constant population sizes (Figure 2). We also considered models similar to IIM2, but assuming unidirectional gene flow from *D. p. bogotana* to *D. persimilis* (IIM3) or from *D. persimilis* to *D. p. bogotana* (IIM4). Nested model comparison supports the choice of any of the three models with varying population sizes (IIM2, IIM3, or IIM4) over IIM1, and the likelihood of IIM2 supports bidirectional gene flow (Table 1). These results provide additional support for gene flow between the *D. persimilis* and *D. pseudoobscura* lineages after speciation and demonstrate that a significant amount of this exchange was relatively ancient (region 2 in Figure 1), occurring prior to the split of *D. p. bogotana.*

### D-statistics suggest recent genetic exchange

Given the evidence for gene flow between *D. persimilis* and *D. pseudoobscura*, we next examined the timing of this gene flow. To contrast sympatric and allopatric subspecies of *D. pseudoobscura* in their similarity to *D. persimilis*, we implemented Patterson’s D-statistic using the tree: (((*D. p. bogotana*, *D. p. pseudoobscura), D. persimilis), D. lowei))).* Patterson’s D-statistic is an implementation of ABBA-BABA, which uses parsimony informative sites to test whether derived alleles (“B”) in *D. persimilis* are shared with *D. p. bogotana* or with *D. p. pseudoobscura* at equal frequencies. Derived alleles in *D. persimilis* may be shared with *D. p. pseudoobscura* due to ancestral polymorphism, ancient gene flow (prior to the split of the two *D. pseudoobscura* subspecies), recent gene flow (since the split of the two *D. pseudoobscura* subspecies), or a combination of these factors. The null expectation is that the two phylogeny-discordant patterns, ABBA and BABA, should be present equally if ancestral polymorphism and ancient gene flow are the sole drivers of patterns of divergence. Gene flow between *D. p. pseudoobscura* and *D. persimilis* since the split of the two *D. pseudoobscura* subspecies (estimated at 150,000 years ago: Schaeffer & Miller 1991) would promote an excess of ABBA over BABA patterns, particularly on freely recombining chromosomes. Indeed, ABBA sites exceed BABA sites on all chromosomes (Table 2), and chromosome 4 shows an unambiguously significant excess of ABBA (|Z-score| >= 5), suggesting that the phylogenetic relationship between these four taxa does not fully explain the observed patterns of divergence and some very recent gene exchange has occurred between the North American species. Furthermore, the genome wide z-score for collinear regions is significant (|Z-score| = 7.215).

**Table 2.** D-statistics. For each region^†^, the number of sites with BABA and ABBA patterns are provided along with the total number of SNPs at sites where all four populations have data.

| Region | D-Statistic | Standard Error | Z-score | BABA Sites | ABBA Sites | SNPs |
| --- | --- | --- | --- | --- | --- | --- |
| *Whole-Genome* |  |  |  |  |  |  |
| Inverted Regions | -0.0111 | 0.0046 | -2.401 | 10,269 | 10,500 | 2,063,794 |
| Collinear Regions | -0.0179 | 0.0020 | -9.154** | 44,162 | 45,776 | 6,180,121 |
| Collinear<sub>FR</sub> Regions | -0.0170 | 0.0024 | -7.215** | 30,749 | 31,813 | 3,750,444 |
| *Chromosome 4* |  |  |  |  |  |  |
| Whole Chromosome | -0.0142 | 0.0027 | -5.335** | 22,606 | 23,256 | 2,526,132 |
| *Chromosome 2* |  |  |  |  |  |  |
| Whole Chromosome | -0.0149 | 0.0032 | -4.713* | 18,654 | 19,220 | 2,833,597 |
| Inverted Region | -0.0044 | 0.0084 | -0.531 | 3,381 | 3,411 | 695,574 |
| Collinear Regions | -0.0172 | 0.0033 | -5.099** | 15,273 | 15,809 | 2,138,123 |
| Collinear<sub>FR</sub> Region | -0.0176 | 0.0048 | -3.686* | 7,061 | 7,314 | 927,665 |
| *Chromosome XL* |  |  |  |  |  |  |
| Whole Chromosome | -0.0161 | 0.0053 | -3.041* | 7,860 | 8,117 | 1,659,395 |
| Inverted Region | -0.0036 | 0.0099 | -0.364 | 2,323 | 2,340 | 606,742 |
| Collinear Regions | -0.0213 | 0.0061 | -3.476* | 5,537 | 5,778 | 1,052,653 |
| Collinear<sub>FR</sub> Region | -0.0388 | 0.0125 | -3.114* | 1,513 | 1,635 | 245,552 |
| *Chromosome XR* |  |  |  |  |  |  |
| Whole Chromosome | -0.0174 | 0.0045 | -3.881* | 11,086 | 11,479 | 1,978,707 |
| Inverted Region | -0.0173 | 0.0066 | -2.633 | 5,236 | 5,420 | 893,006 |
| Collinear Region | -0.0176 | 0.0061 | -2.875 | 5,849 | 6,059 | 1,085,701 |
| Collinear<sub>FR</sub> Region | -0.0224 | 0.0133 | -1.688 | 1,577 | 1,649 | 265,259 |

<sup>†</sup> The presented chromosomal regions include entire chromosomes, inverted regions only, collinear regions only, and collinear regions excluding sequence within 5 mb of chromosome ends or within 2.5 mb of inversion breakpoints (Collinear<sub>FR</sub>).
\* Significant at $|z| \geq 3$ .
\*\* Significant at $|z| \geq 5$ .

Given the observed excess of ABBA over BABA sites throughout the genome, we next applied f_d_ to quantify this excess in smaller genomic intervals. In comparison to D-statistics, f_d_ is less affected by differences in effective population size and is better suited to identifying introgression regions (Martin *et al.* 2015). The genome-wide patterns of f_d_ support the evidence of gene flow between *D. p. pseudoobscura* and *D. persimilis*, particularly in the collinear regions of the genome (Figure 4). Inverted regions exhibit markedly lower f_d_ compared to collinear regions (Figure 4A). This difference is statistically significant on all inversion-bearing chromosomes, regardless of whether the inverted regions are compared to all collinear regions or just the conservative subset contained in collinear_FR_ (Figure 4B; p < 0.01 for all comparisons, Mann-Whitney U).

**Figure 4.**
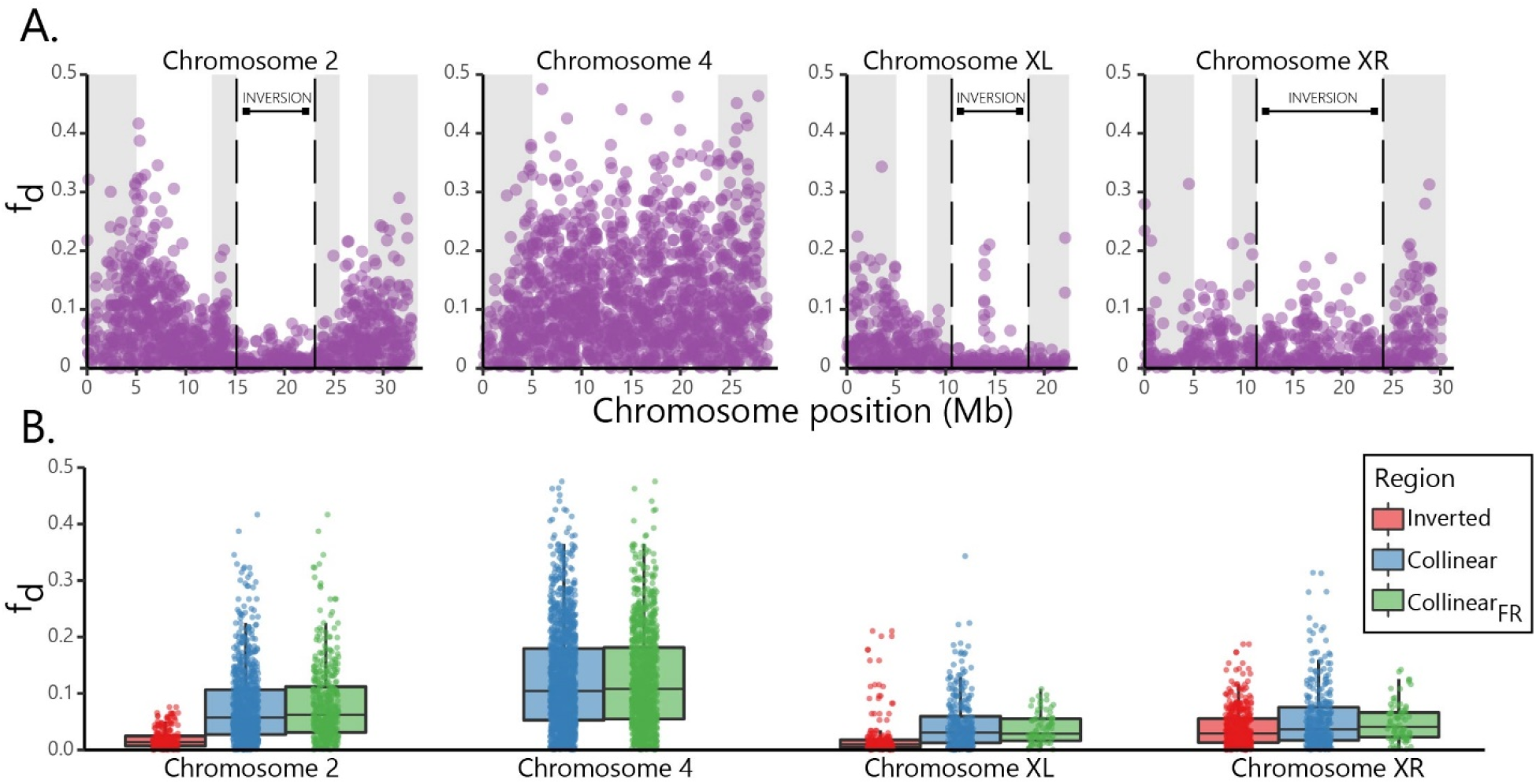
Signals of introgression along the genome. (A) The estimated proportion of introgression (f_d_) between *D. pseudoobscura* and *D. persimilis* is shown in non-overlapping 100 SNP windows along chromosomes 2, 4, XL, and XR. Inversion boundaries are shown with dashed black lines, and collinear regions are grayed-out where they approach inversion breakpoints or chromosome ends (windows excluded from Collinear_FR_). (B) Summarizes these introgression estimates by region: Inverted, Collinear, and Collinear_FR_.

### Is recent gene flow responsible for patterns of higher divergence in allopatry vs sympatry?

All three inversion-bearing chromosomes exhibit lower divergence in the *D. persimilis:D. p. pseudoobscura* comparison vs the *D. persimilis:D. p. bogotana* comparison (Figure 3). Notably, this lower divergence in the sympatric comparison is statistically significant for both the collinear and inverted regions (p < 0.001 on each chromosome, Mann-Whitney U test). Divergence between the species pairs in both inverted and collinear regions shows the magnitude of this difference (Figure 3). This pattern is consistent across all strains: differentiation in the inverted regions between *D. persimilis:D. p. bogotana* is higher than differentiation between *D. persimilis* and any of the North American *D. p. pseudoobscura* genomes (Supplementary Figure 2).

The observation that divergence is lower in sympatry compared to allopatry even in the inverted regions is somewhat surprising. A possible explanation for the lower divergence in the sympatric species pair is that gene flow is homogenizing these species even in inverted regions. Though there is evidence that double crossovers and gene conversions occur within inversions (Schaeffer & Anderson 2005; Stevison *et al.* 2011; Crown *et al.* 2018; Korunes & Noor 2019), an alternative explanation is that *D. p. bogotana* has experienced more substitutions per site, possibly due to differences in demographic history among the lineages. *D. p. bogotana* may have experienced a population bottleneck upon colonization of South America leading to a subsequently small effective population size (Schaeffer & Miller 1991; Wang & Hey 1996; Machado *et al.* 2002), which might allow drift to result in a higher fixation rate of slightly deleterious mutations (Whitlock 2000; Charlesworth 2009). Indeed, genome-wide comparison of the relative substitution rates (Tajima 1993) between the lineages reveals that *D. p. bogotana* has experienced significantly more substitutions per site than *D. p. pseudoobscura* relative to the outgroup species, *D. lowei* (Supplementary Table 6)

To correct for differences in evolutionary rates in these populations, we considered the effects of introgression on divergence after adjusting for an elevated rate of fixation leading to a longer branch length in *D. p. bogotana.* We compared the divergence of each *D. pseudoobscura* subspecies from *D. persimilis* to the divergence of each *D. pseudoobscura* subspecies from *D. lowei* using the following equation to define the “introgression effect”: (D_xy_ [*D.persimilis*:*D.p.bogotana*] − D_xy_ [*D.persimilis*:*D.p.pseudo.*]) − (D_xy_ [*D.lowei*:*D.p.bogotana*] − D_xy_ [*D.lowei*:*D.p.pseudo*.]). The first half of this equation should include the effects of branch length in *D. p. bogotana* and the effects of any introgression between *D. pseudoobscura* and *D. persimilis* (Figure 5A). Since *D. lowei* does not hybridize with any of these species, the second half of the equation should reflect only the effects of branch length in *D. p. bogotana*. Thus, the difference between these terms should subtract the effects of evolutionary rate, leaving the effects of recent introgression. As we are interested in whether, when, and how chromosomal inversions are contributing to patterns of divergence by suppressing gene flow, we compared inverted vs. collinear regions in their introgression effects. After subtracting the effects of branch length in *D. p. bogotana*, there is still evidence that differential gene flow in inverted vs collinear regions has statistically significantly influenced patterns of divergence between *D. pseudoobscura* and *D. persimilis.* The distributions of the introgression effect values in each of the sampled *D. p. pseudoobscura* genomes differs significantly between inverted and collinear regions (p < 0.001, Mann-Whitney U test; Figure 5B). Notably, the introgression effect is positive, albeit small, in the inverted regions, potentially due to double crossovers and gene conversions occurring within inversions (Schaeffer & Anderson 2005; Stevison *et al.* 2011; Crown *et al.* 2018; Korunes & Noor 2019).

**Figure 5.**
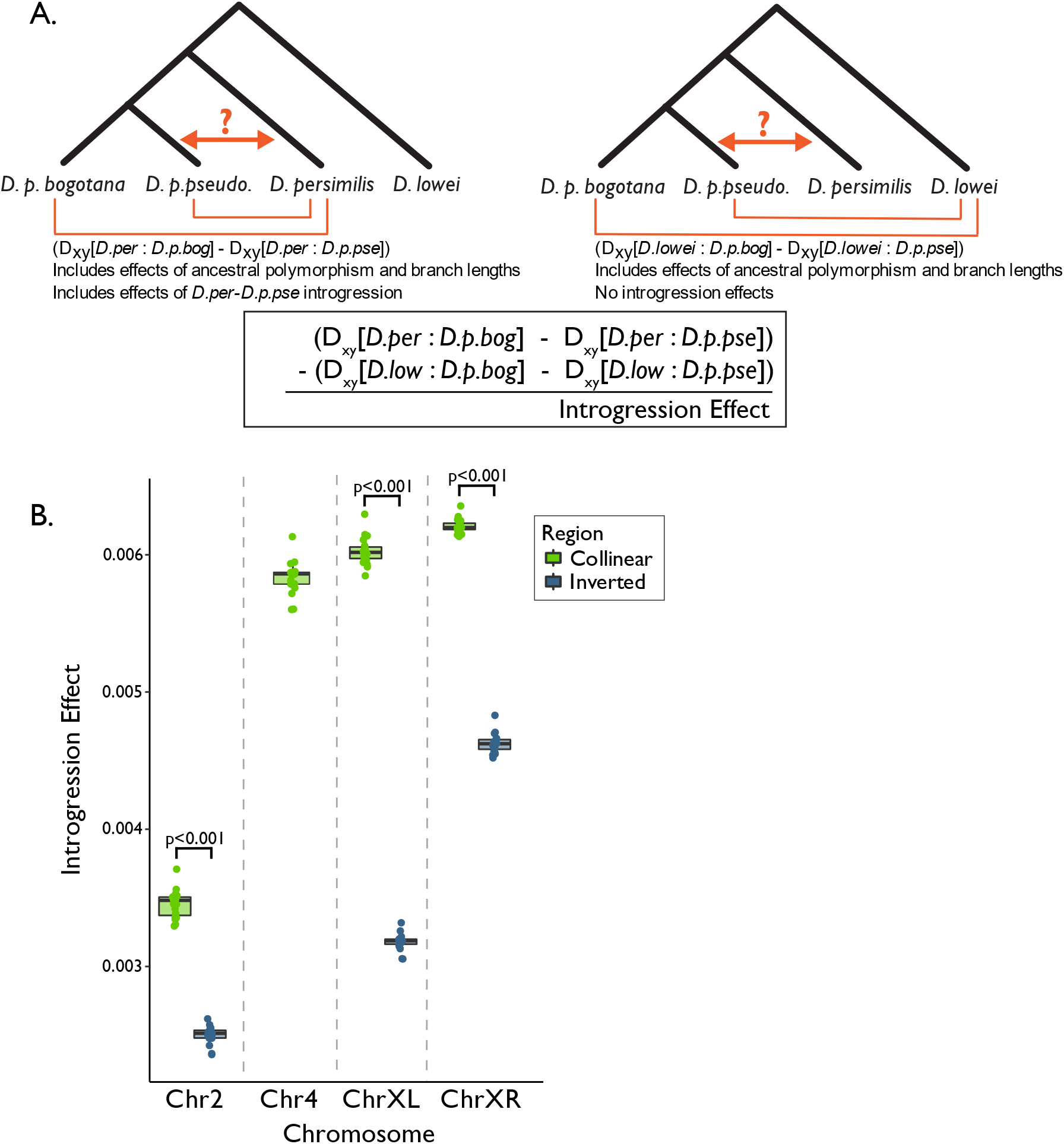
The effects of introgression on divergence after correcting for evolutionary rate. (A) Shows the strategy for examining the “introgression effect” of gene flow on divergence. The first half of the equation compares the divergence of each *D. pseudoobscura* subspecies from *D. persimilis*, and the second half compares the divergence of each *D. pseudoobscura* subspecies from *D. lowei.* The difference between these terms, plotted in (B) for each of the sampled *D. p. pseudoobscura* genomes, should reflect the effects of introgression after correcting for branch length.

## Discussion

Our model-based examination of gene flow and our application of Patterson’s D-statistic to examine shared variation found between *D. persimilis* and sympatric and allopatric subspecies of *D. pseudoobscura* indicates post-speciation gene exchange, including both early post-speciation gene flow and gene flow within the past ~150,000 years. Specifically, we interpret the excess of ABBA > BABA sites as evidence that *D. pseudoobscura* and *D. persimilis* have exchanged genes in collinear regions since the split of *D. p. bogotana*. Our analyses span the majority of the assembled genome, excluding only the ~17% of the genome where we cannot reasonably apply these methods to examine introgression and divergence (see *Methods*).

This evidence that introgression is driving patterns of divergence between *D. pseudoobscura* and *D. persimilis* is in agreement with previous reports of ongoing hybridization in these species (Dobzhansky 1973; Powell 1983; Machado & Hey 2003; Hey & Nielsen 2004). Despite our evidence for recent gene exchange, it appears that introgression is not the sole driver of patterns of divergence between these species overall. While D-statistics and f_d_ suggest an excess of shared, derived alleles across the genomes of *D. pseudoobscura* and *D. persimilis*, these statistics may be biased by factors such as ancestral population structure and differences in effective population size (Slatkin & Pollack 2008; Martin *et al.* 2015; He *et al.* 2020). In comparison to Patterson’s D-statistic, f_d_ is less sensitive to local variation in recombination rate and divergence. However, it can still be biased by regions of reduced interspecies divergence, which may distort tests for recent introgression (Martin *et al.* 2015), so the conclusion of recent introgression would be tentative based on these results alone. Here, we explore other important factors that might underlie the observed patterns of divergence, with particular consideration of how these factors might confound signals of recent introgression.

As seen in Figure 3 and in previous studies (Kulathinal *et al.* 2009), divergence is higher in inverted vs collinear regions in this system. This difference holds for divergence between *D. persimilis* and either *D. p. bogotana* or *D. p. pseudoobscura*. The lower observed divergence in sympatry compared to allopatry even in the inverted regions is somewhat surprising given the expectation that recombination in hybrids will be restricted in these inverted regions. This observation led us to consider the possibility that the allopatric subspecies, *D. p. bogotana*, might have experienced more nucleotide substitutions per site than the other taxa. Thus, we considered four non-mutually exclusive factors that might contribute to the observed patterns of divergence with respect to chromosomal arrangement: 1) the segregation of ancestral polymorphism (as advocated by Fuller *et al*. (2018)), 2) increased branch length in the allopatric *D. p. bogotana*, 3) gene flow prior to the split of *D. p. bogotana*, and 4) recent/ongoing gene flow (the latter two discussed in Powell 1983; Wang & Hey 1996; Wang *et al.* 1997; Noor *et al.* 2001, 2007; Machado & Hey 2003; Hey & Nielsen 2004; Machado *et al.* 2007; Kulathinal *et al.* 2009; McGaugh & Noor 2012). Achieving a cohesive view of the role of inversions in species divergence relies on considering the combined effects of these factors.

For the sympatric pair *D. persimilis* vs *D. p. pseudoobscura*, any difference in D_xy_ in inverted regions compared to collinear regions could be due to the segregation of ancestral inversion polymorphism or to post-speciation genetic exchange. In contrast, any difference in D_xy_ in inverted regions compared to collinear regions in the allopatric pair *D. persimilis* vs *D. p. bogotana* could be driven by the segregation of ancestral inversion polymorphism or by post-speciation gene flow prior to the split of *D. p. bogotana.* This comparison will not reflect any recent gene flow, since *D. p. bogotana* has evolved in allopatry for the past 150,000 years. We leveraged this contrast to isolate the effects of recent introgression on divergence, and our findings here suggest that contributions from recent gene flow only partially explain observed divergence patterns.

The observed "introgression effect" can be thought of as an indicator of the relative influence of recent introgression on the reduction in *D. persimilis* vs *D. pseudoobscura* divergence in sympatry vs allopatry. Our method does not account for all evolutionary forces shaping nucleotide divergence, so we do not consider it a quantitative measurement of introgression. However, it provides a useful way to consider the relative contribution of recent introgression compared to ancestral polymorphism and branch length. The second half of the introgression effect equation (Figure 5A; D_xy_[*D.lowei*:*D.p.bogotana*] − D_xy_[*D.lowei*:*D.p.pseudo*.]) includes the effects of branch length in *D. p. bogotana* and the effects of ancestral polymorphism without including any effects of recent introgression. We can subtract this term from D_xy_[*D.persimilis:D.p.bogotana*] to obtain a hypothetical D_xy_ reflecting what divergence between *D. persimilis* and *D. p. pseudoobscura* might look like without the effects of recent introgression, ancestral polymorphism, or longer branch length in *D. p. bogotana*. Dividing the introgression effect by this hypothetical D_xy_ gives the hypothetical change in D_xy_ in allopatry vs sympatry due specifically to recent introgression. Given the observed average introgression effect (0.0045) and the hypothetical D_xy_[*D. persimilis*:*D. p. pseudoobscura*] of 0.01, recent introgression might explain roughly half of the reduction in D_xy_ observed in sympatry vs allopatry. As such, the homogenization effect of recent introgression on sequence divergence appears far from trivial.

These results suggest that patterns of divergence between *D. persimilis* and *D. pseudoobscura* are explained by a combination of segregating ancestral polymorphism and post-speciation gene flow. We applied a model-based approach to investigate the timing of gene flow between *D. persimilis* and *D. pseudoobscura.* Our results suggest that an isolation-with-initial-migration model best explains the divergence of *D. persimilis* and *D. p. bogotana* when compared to a model of strict isolation. This result provides further evidence for gene flow between *D. persimilis* and *D. pseudoobscura*, and it suggests that some of this gene flow occurred prior to the split of *D. p. bogotana* and remains detectable in observed genetic patterns.

Our results question interpretations from earlier studies of this system. Given that *D. p. bogotana* can be reasonably assumed to not be currently exchanging genes with either *D. persimilis* or *D. p. pseudoobscura* (Schaeffer & Miller 1991; Wang *et al.* 1997), *D. persimilis*:*D. p. bogotana* divergence was argued to be a suitable “negative control” for examining the effect of recent hybridization between *D. persimilis* and *D. p. pseudoobscura* (Brown *et al.* 2004). By this argument, the effect of recent gene flow can be estimated by an allopatric vs sympatric comparison of the difference in divergence (whether in DNA sequence or in phenotype) in inverted regions to divergence in collinear regions. Specifically, Brown *et al*. (2004) and Chang and Noor (2007) inferred multiple hybrid sterility factors between *D. p. bogotana* and *D. persimilis* that did not distinguish North American *D. p. pseudoobscura* and *D. persimilis* (Brown *et al.* 2004; Chang & Noor 2007). Similarly, Kulathinal *et al.* (2009) observed significantly greater sequence difference between *D. p. bogotana* and *D. persimilis* than between *D. p. pseudoobscura* and *D. persimilis*. In both cases, the authors interpreted the difference to result from recent homogenization of the collinear regions in the latter pair. Based on our findings, we suggest this difference may result at least in part from the accelerated rate of divergence in *D. p. bogotana*.

Overall, we caution that simple allopatry-sympatry comparisons can easily be misleading, and the population histories and rates of evolution of the examined species should be carefully considered. Variation in N_e_ due to events such as recent population bottlenecks in one of the taxa can dramatically influence evolutionary rates (reviewed in Charlesworth 2009; Lanfear *et al.* 2014). Additionally, the process of genetic divergence that shapes alleles responsible for local adaptation and hybrid incompatibility can extend deep into the history of the species. In fact, the influence of inversions on the divergence of a species pair can predate the split of the species. Inversion polymorphisms in the ancestral population of a species pair can contribute to patterns of higher sequence differentiation between species in those inverted regions (Fuller *et al.* 2018). Separating these effects requires an understanding of the timing and extent of introgression, which can only be understood with an appreciation for the evolutionary processes occurring in each of the taxa at hand. Though there are many remaining questions about how inversions shape divergence, we present evidence that inversions have contributed to the divergence of *D. pseudoobscura* and *D. persimilis* over multiple distinct periods during their speciation: 1) pre-speciation segregation of inversions in the ancestral population, 2) post-speciation gene flow prior to the split of *D. p. bogotana*, and 3) recent gene flow.

## Acknowledgements

We thank all members of the Noor lab for helpful discussions, and we thank Zachary Fuller and Russell Corbett-Detig for their thoughtful feedback on the manuscript. KLK was supported by the National Science Foundation Graduate Research Fellowship Program under Grant No. DGE 1644868, and this work was additionally supported by National Science Foundation grants DEB-1754022 and DEB-1754439 to MAFN, and MCB-1716532 and DEB-1754572 to CAM.

## References

Altschul SF, Gish W, Miller W, Myers EW, Lipman DJ (1990) Basic local alignment search tool. Journal of Molecular Biology, 215, 403–410.

Andolfatto P, Wall JD (2003) Linkage disequilibrium patterns across a recombination gradient in African Drosophila melanogaster. Genetics, 165, 1289–1305.

Van der Auwera GA, Carneiro MO, Hartl C et al. (2013) From FastQ data to high-confidence variant calls: The Genome Analysis Toolkit best practices pipeline. In: Current Protocols in Bioinformatics, pp. 11.10.1–11.10.33. John Wiley & Sons, Inc., Hoboken, NJ, USA.

Ayala FJ, Coluzzi M (2005) Chromosome speciation: humans, Drosophila, and mosquitoes. Proceedings of the National Academy of Sciences, 102 Suppl, 6535–42.

Beckenbach AT, Wei YW, Liu H (1993) Relationships in the Drosophila obscura species group, inferred from mitochondrial cytochrome oxidase II sequences. Molecular Biology and Evolution, 10, 619–634.

Brown KM, Burk LM, Henagan LM, Noor MAF (2004) A test of the chromosomal rearrangement model of speciation in Drosophila pseudoobscura. Evolution, 58, 1856–60.

Butlin RK (2005) Recombination and speciation. Molecular Ecology, 14, 2621–2635.

Chang AS, Noor MAF (2007) The genetics of hybrid male sterility between the allopatric species pair Drosophila persimilis and D. pseudoobscura bogotana: dominant sterility alleles in collinear autosomal regions. Genetics, 176, 343–9.

Charlesworth B (2009) Effective population size and patterns of molecular evolution and variation. Nature Reviews Genetics, 10, 195–205.

Costa RJ, Wilkinson-Herbots H (2017) Inference of gene flow in the process of speciation: An efficient maximum-likelihood method for the isolation-with-initial-migration model. Genetics, 205, 1597–1618.

Crown KN, Miller DE, Sekelsky J, Hawley RS (2018) Local inversion heterozygosity alters recombination throughout the genome. Current Biology, 28, 2984–2990.

Dobzhansky T (1973) Is there gene exchange between Drosophila pseudoobsura and Drosophila persimilis in their natural habitats? The American Naturalist, Vol. 107, No. 954, pp. 312–314.

Dobzhansky T, Epling C (1944) Contributions to the genetics, taxonomy, and ecology of Drosophila pseudoobscura and its relatives. Carnegie Institute of Washington, 544, 1–46.

Feder JL, Gejji R, Powell THQ, Nosil P (2011) Adaptive chromosomal divergence driven by mixed geographic mode of evolution. Evolution, 65, 2157–2170.

Fuller ZL, Leonard CJ, Young RE, Schaeffer SW, Phadnis N (2018) Ancestral polymorphisms explain the role of chromosomal inversions in speciation. PLoS Genetics, 14, e1007526.

Guerrero RF, Rousset F, Kirkpatrick M (2012) Coalescent patterns for chromosomal inversions in divergent populations. Philosophical transactions of the Royal Society B, 367, 430–8.

He C, Liang D, Zhang P (2020) Asymmetric Distribution of Gene Trees Can Arise under Purifying Selection If Differences in Population Size Exist. Molecular biology and evolution, 37, 881–892.

Heed WB, Crumpacker DW, Ehrman L (1969) Drosophila lowei, a new American member of the Obscura species group1,2. Annals of the Entomological Society of America, 62, 388–393.

Hey J, Nielsen R (2004) Multilocus methods for estimating population sizes, migration rates and divergence time, with applications to the divergence of Drosophila pseudoobscura and D. persimilis. Genetics, 167, 747–60.

Jackson BC (2011) Recombination-suppression: how many mechanisms for chromosomal speciation? Genetica, 139, 393–402.

Korunes KL, Noor MAF (2019) Pervasive gene conversion in chromosomal inversion heterozygotes. Molecular Ecology, 28, 1302–1315.

Kulathinal RJ, Bennett SM, Fitzpatrick CL, Noor MAF (2008) Fine-scale mapping of recombination rate in Drosophila refines its correlation to diversity and divergence. Proceedings of the National Academy of Sciences, 105, 10051–6.

Kulathinal RJ, Stevison LS, Noor MAF (2009) The genomics of speciation in Drosophila: diversity, divergence, and introgression estimated using low-coverage genome sequencing. PLoS Genetics, 5, e1000550.

Lanfear R, Kokko H, Eyre-Walker A (2014) Population size and the rate of evolution. Trends in Ecology and Evolution, 29, 33–41.

Langley CH, Lazzaro BP, Phillips W, Heikkinen E, Braverman JM (2000) Linkage disequilibria and the site frequency spectra in the su(s) and su(wa) regions of the Drosophila melanogaster X chromosome. Genetics, 156, 1837–1852.

Li H, Durbin R (2009) Fast and accurate short read alignment with Burrows-Wheeler Transform. Bioinformatics, 25, 1754–60.

Li H, Handsaker B, Wysoker A et al. (2009) The Sequence Alignment/Map format and SAMtools. Bioinformatics, 25, 2078–9.

Lowry DB, Willis JH (2010) A widespread chromosomal inversion polymorphism contributes to a major life-history transition, local adaptation, and reproductive isolation. PLoS Biology, 8, e1000500.

Machado CA, Haselkorn TS, Noor MAF (2007) Evaluation of the genomic extent of effects of fixed inversion differences on intraspecific variation and interspecific gene flow in Drosophila pseudoobscura and D. persimilis. Genetics, 175, 1289–306.

Machado CA, Hey J (2003) The causes of phylogenetic conflict in a classic Drosophila species group. Proceedings of the Royal Society B: Biological Sciences, 270, 1193–202.

Machado CA, Kliman RM, Markert JA, Hey J (2002) Inferring the history of speciation from multilocus DNA sequence data: The case of Drosophila pseudoobscura and close relatives. Molecular Biology and Evolution, 19, 472–488.

Malinsky M, Matschiner M, Svardal H (2020) Dsuite - fast D-statistics and related admixture evidence from VCF files. bioRxiv.

Mallet J (2005) Hybridization as an invasion of the genome. Trends in ecology & evolution, 20, 229–37.

Manoukis NC, Powell JR, Touré MB et al. (2008) A test of the chromosomal theory of ecotypic speciation in Anopheles gambiae. Proceedings of the National Academy of Sciences of the United States of America, 105, 2940–2945.

Martin SH, Davey JW, Jiggins CD (2015) Evaluating the use of ABBA–BABA statistics to locate introgressed loci. Molecular Biology and Evolution, 32, 244–257.

McGaugh SE, Noor MAF (2012) Genomic impacts of chromosomal inversions in parapatric Drosophila species. Philosophical Transactions of the Royal Society B: Biological Sciences, 367, 422–9.

McKenna A, Hanna M, Banks E et al. (2010) The Genome Analysis Toolkit: a MapReduce framework for analyzing next-generation DNA sequencing data. Genome Research, 20, 1297–303.

Michel AP, Sim S, Powell THQ et al. (2010) Widespread genomic divergence during sympatric speciation. Proceedings of the National Academy of Sciences of the United States of America, 107, 9724–9729.

Noor MAF, Garfield DA, Schaeffer SW, Machado CA (2007) Divergence between the Drosophila pseudoobscura and D. persimilis genome sequences in relation to chromosomal inversions. Genetics, 177, 1417–28.

Noor MA, Grams KL, Bertucci LA, Reiland J (2001) Chromosomal inversions and the reproductive isolation of species. Proceedings of the National Academy of Sciences, 98, 12084–8.

Patterson N, Moorjani P, Luo Y et al. (2012) Ancient admixture in human history. Genetics, 192, 1065–93.

Payseur BA, Rieseberg LH (2016) A genomic perspective on hybridization and speciation. Molecular ecology, 25, 2337–60.

Powell JR (1983) Interspecific cytoplasmic gene flow in the absence of nuclear gene flow: Evidence from Drosophila. Proceedings of the National Academy of Sciences, 80, 492–5.

Powell JR (1992) Inversion polymorphisms in Drosophila pseudoobscura and Drosophila persimilis. Drosophila Inversion Polymorphism, 73–126.

Purcell S, Neale B, Todd-Brown K et al. (2007) PLINK: a tool set for whole-genome association and population-based linkage analyses. American journal of human genetics, 81, 559–75.

Schaeffer SW, Anderson WW (2005) Mechanisms of genetic exchange within the chromosomal inversions of Drosophila pseudoobscura. Genetics, 171, 1729–39.

Schaeffer SW, Bhutkar A, McAllister BF et al. (2008) Polytene chromosomal maps of 11 Drosophila species: The order of genomic scaffolds inferred from genetic and physical maps. Genetics, 179, 1601–55.

Schaeffer SW, Miller EL (1991) Nucleotide sequence analysis of Adh genes estimates the time of geographic isolation of the Bogota population of Drosophila pseudoobscura. Proceedings of the National Academy of Sciences, 88, 6097–101.

Slatkin M, Pollack JL (2008) Subdivision in an ancestral species creates asymmetry in gene trees. Molecular biology and evolution, 25, 2241–6.

Stevison LS, Hoehn KB, Noor MAF (2011) Effects of inversions on within- and between-species recombination and divergence. Genome Biology and Evolution, 3, 830–41.

Stevison LS, Noor MAF (2010) Genetic and evolutionary correlates of fine-scale recombination rate variation in Drosophila persimilis. Journal of Molecular Evolution, 71, 332–345.

Tajima F (1993) Simple methods for testing the molecular evolutionary clock hypothesis. Genetics, 135.

Taylor SA, Larson EL (2019) Insights from genomes into the evolutionary importance and prevalence of hybridization in nature. Nature Ecology and Evolution, 3, 170–177.

Wang RL, Hey J (1996) The speciation history of Drosophila pseudoobscura and close relatives: inferences from DNA sequence variation at the period locus. Genetics, 144, 1113–26.

Wang Y, Hey J (2010) Estimating divergence parameters with small samples from a large number of loci. Genetics, 184, 363–379.

Wang RL, Wakeley J, Hey J (1997) Gene flow and natural selection in the origin of Drosophila pseudoobscura and close relatives. Genetics, 147, 1091–1106.

Whitlock MC (2000) Fixation of new alleles and the extinction of small populations: Drift load, beneficial alleles, and sexual selection. Evolution, 54, 1855–1861.

Yang Z (2002) Likelihood and Bayes estimation of ancestral population sizes in hominoids using data from multiple loci. Genetics, 162, 1811–23.

