## Supplementary Figures and Tables for "Inversions shape the divergence of *Drosophila pseudoobscura* and *D. persimilis* on multiple timescales"

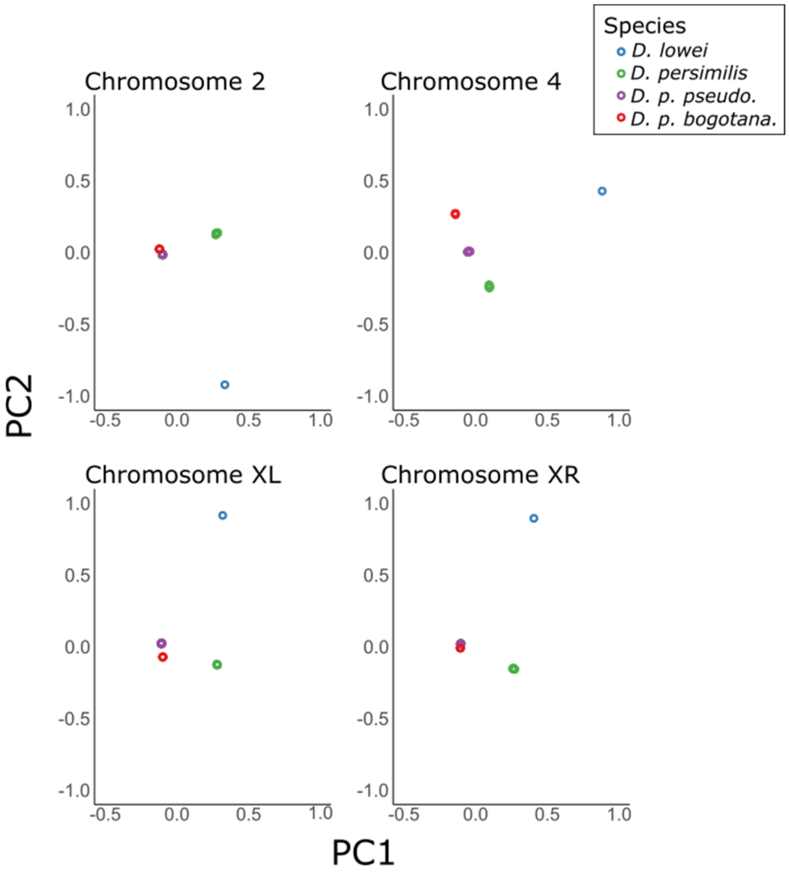


**Supplementary Figure 1** | Principal components analysis. The first two principal components are plotted for the four examined chromosome arms.

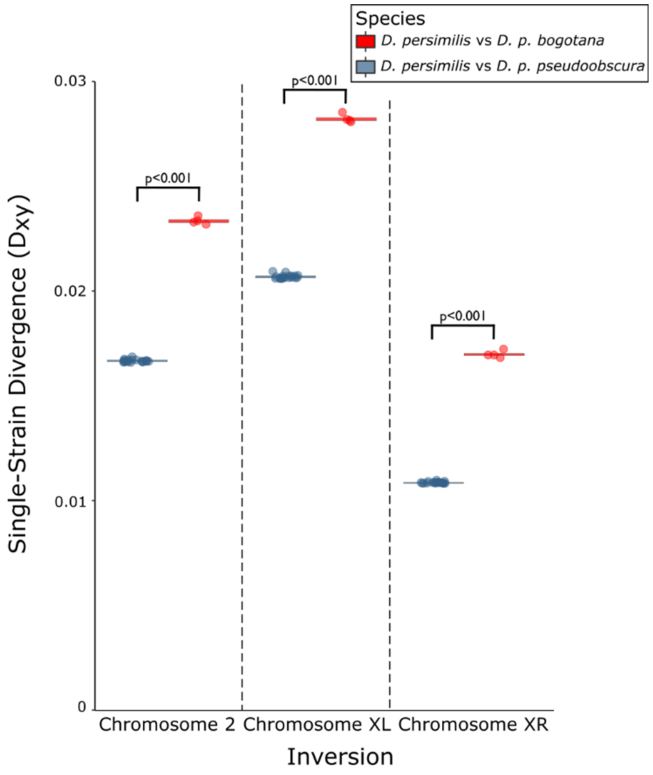


**Supplementary Figure 2** | Divergence in inverted regions. For each individual *D. pseudoobscura* genome, divergence from *D. persimilis* is shown for inverted regions.

**Supplementary Table 1** | Genomic data for introgression analysis.

| Strains | Sequence Accessions |
|---|---|
| ***Drosophila lowei*** | |
| 1 strain: Lab3Lowei | Raw sequences available under SRA Experiment SRX091467. |
| ***Drosophila persimilis*** | |
| 8 strains: SCI, MSH7, MSH42, MatherG, Mather40, 111_51, 111_50, and 111_35 | Raw sequences will be published to SRA by the time of publication. |
| ***Drosophila pseudoobscura bogotana*** | |
| 4 strains: Toro4, Suta3, ER-White, SCinv | Raw sequences available under SRA Project PRJNA593268. |
| ***Drosophila pseudoobscura pseudoobscura*** | |
| 19 strains: S1-A56, S10-A47, S11-A14, S12-M27, S13-A48, S14-A49, S15-A57, S16-A30, S17-M20, S18-M15, S19-A24, S20-M13, S21-M6, S22-A6, S3-M14, S4-A60, S5-M17, S6-A19, S9-A12 | Raw sequences will be published to SRA by the time of publication. |

**Supplementary Table 2** | Inversion breakpoints relative to the *D. miranda* reference genome. Breakpoints (Chr 2, XL, XR) were originally described in Machado *et al.* (2007) and provided here in coordinates of DroMir2.2.

| Chromosome | Approximate Proximal Breakpoint | Approximate Distal Breakpoint |
|---|---|---|
| Chromosome 2 | 15,154,864 | 23,081,317 |
| Chromosome XL | 10,619,782 | 18,389,958 |
| Chromosome XR | 24,174,893 | 11,348,640 |

**Supplementary Table 3** | Average divergence in inverted regions compared to collinear regions[1].

| Chromosome | Average $D_{xy:}$ Inverted | Average $D_{xy}$ Collinear | Average $D_{xy}$ $Collinear_{FR}$ | p-value Inverted vs Collinear | p-value Inverted vs $Collinear_{FR}$ |
|---|---|---|---|---|---|
| ***D. persimilis* vs *D. p. pseudoobscura*** | | | | | |
| Chromosome 2 | 0.0166 | 0.0125 | 0.0131 | <0.001 | <0.001 |
| Chromosome XL | 0.0207 | 0.0143 | 0.0130 | <0.001 | <0.001 |
| Chromosome XR | 0.0109 | 0.0094 | 0.0089 | <0.001 | <0.001 |
| ***D. persimilis* vs *D. p. bogotana*** | | | | | |
| Chromosome 2 | 0.0233 | 0.0198 | 0.0203 | <0.001 | <0.001 |
| Chromosome XL | 0.0283 | 0.0235 | 0.0238 | <0.001 | <0.001 |
| Chromosome XR | 0.0171 | 0.0168 | 0.0153 | 0.202 | <0.001 |

[1] Statistical comparisons of these estimates were made using Mann-Whitney U tests to compare the divergence estimates in each region, split into 50 kb windows.

**Supplementary Table 4** | Using loci sampled from $collinear_{FR}$ regions only, forward selection of the best model of *D. persimilis - D. p. bogotana* divergence using the maximized log-likelihood (LogL) under each model in likelihood-ratio tests.

| $H_0$ | $H_1$ | Deg. of Freedom | LogL $H_0$ | LogL $H_1$ | LRT Statistic | P-value |
|---|---|---|---|---|---|---|
| Iso | IM | 2 | -25511.23 | -25492.84 | 36.78 | 1.031e-08 |
| IM | IIM1 | 1 | -25492.84 | -25492.84 | 0 | - |
| IM | IIM2 | 3 | -25492.84 | -25442.42 | 100.84 | 1.025e-21 |
| IIM1 | IIM2 | 2 | -25492.84 | -25442.42 | 100.84 | 1.267e-22 |
| IIM1 | IIM3 | 1 | -25492.84 | -25447.73 | 90.22 | 2.131e-21 |
| IIM1 | IIM4 | 1 | -25492.84 | -25444.03 | 97.62 | 5.069e-23 |

**Supplementary Table 5** | Maximum-likelihood estimates under models of *D. persimilis* - *D. pseudoobscura bogotana* divergence and values of the maximized log-likelihood (LogL).

| Model | $\theta_1$ | $\theta_2$ | $\theta_A$ | $\theta_{C1}$ | $\theta_{C2}$ | V | T1 | M1 | M2 | logL |
|---|---|---|---|---|---|---|---|---|---|---|
| *Loci sampled genome-wide* | | | | | | | | | | |
| **Divergence in Isolation** | | | | | | | | | | |
| ISO | 2.756 | 1.992 | 3.468 | - | - | 3.626 | - | - | - | -33702.73 |
| **Divergence in isolation with migration** | | | | | | | | | | |
| IM | 2.599 | 1.721 | 3.391 | - | - | 4.083 | - | 0 | 0.251 | -33659.39 |
| **Divergence in isolation with initial migration** | | | | | | | | | | |
| IIM1 | 2.599 | 1.721 | 3.391 | - | - | 4.082 | 1e-06 | 0 | 0.251 | -33659.39 |
| IIM2 | 2.631 | 1.950 | 1.825 | 9.587 | 1.830 | 3.380 | 1.089 | 0.187 | 0.164 | -33538.92 |
| IIM3 | 2.661 | 1.715 | 2.303 | 10.453 | 1.839 | 3.413 | 0.909 | - | 0.353 | -33543.76 |
| IIM4 | 2.654 | 2.288 | 1.543 | 9.735 | 1.807 | 3.215 | 1.158 | 0.300 | - | -33542.89 |
| *Loci sampled from collinear regions only* | | | | | | | | | | |
| **Divergence in Isolation** | | | | | | | | | | |
| ISO | 2.857 | 2.026 | 4.120 | - | - | 3.162 | - | - | - | -25511.23 |
| **Divergence in isolation with migration** | | | | | | | | | | |
| IM | 2.724 | 1.738 | 4.082 | - | - | 3.538 | - | 0 | 0.368 | -25492.84 |
| **Divergence in isolation with initial migration** | | | | | | | | | | |
| IIM1 | 2.724 | 1.739 | 4.082 | - | - | 3.533 | 0.005 | 0 | 0.369 | -25492.84 |
| IIM2 | 2.710 | 1.988 | 1.995 | 8.328 | 1.903 | 3.938 | 1.202 | 0.424 | 0.237 | -25442.42 |
| IIM3 | 2.751 | 1.586 | 2.835 | 8.878 | 1.935 | 2.930 | 0.936 | - | 0.694 | -25447.73 |
| IIM4 | 2.732 | 2.371 | 1.718 | 8.616 | 1.873 | 2.820 | 1.234 | 0.569 | - | -25444.03 |

**Supplementary Table 6** | Tajima's relative rate test. Genome-wide comparison of the substitution rates of each lineage compared to the outgroup *D. lowei* (including both inverted/collinear and intergenic/genic SNPs from chromosomes 2, 3, XR, and XL)

| **Genomes Compared to Outgroup (*D. lowei*)** | | **Substitutions** | | **Test Statistic** |
|---|---|---|---|---|
| **Line1** | **Line2** | **$m_1$** (Line1,Outgroup) | **m2** (Line2,Outgroup) | $X^2$ |
| *D. p. bogotana* Line SCinv | *D. p. bogotana* Line Suta3 | 135464 | 135233 | 0.20 |
| *D. p. bogotana* Line SCinv | *D. p. bogotana* Line Toro4 | 133199 | 132939 | 0.25 |
| *D. p. bogotana* Line SCinv | *D. p. bogotana* Line WhiteER | 137670 | 137815 | 0.08 |
| *D. p. bogotana* Line SCinv | *D. pseudoobscura* Line S1-A56 | 362782 | 307843 | 4500.72** |
| *D. p. bogotana* Line SCinv | *D. pseudoobscura* Line S10-A47 | 363835 | 313950 | 3671.54** |
| *D. p. bogotana* Line SCinv | *D. pseudoobscura* Line S11-A14 | 361134 | 302789 | 5127.31** |
| *D. p. bogotana* Line SCinv | *D. pseudoobscura* Line S12-M27 | 355206 | 284279 | 7866.70** |
| *D. p. bogotana* Line SCinv | *D. pseudoobscura* Line S13-A48 | 355405 | 289593 | 6715.09** |
| *D. p. bogotana* Line SCinv | *D. pseudoobscura* Line S14-A49 | 360900 | 304058 | 4858.97** |
| *D. p. bogotana* Line SCinv | *D. pseudoobscura* Line S15-A57 | 363020 | 310913 | 4028.80** |
| *D. p. bogotana* Line SCinv | *D. pseudoobscura* Line S16-A30 | 353165 | 279822 | 8498.11** |
| *D. p. bogotana* Line SCinv | *D. pseudoobscura* Line S17-M20 | 315070 | 151071 | 57698.58** |
| *D. p. bogotana* Line SCinv | *D. pseudoobscura* Line S18-M15 | 353368 | 276725 | 9322.67** |
| *D. p. bogotana* Line SCinv | *D. pseudoobscura* Line S19-A24 | 354527 | 281630 | 8353.24** |
| *D. p. bogotana* Line SCinv | *D. pseudoobscura* Line S20-M13 | 348004 | 256521 | 13844.16** |
| *D. p. bogotana* Line SCinv | *D. pseudoobscura* Line S21-M6 | 354757 | 280638 | 8646.00** |
| *D. p. bogotana* Line Suta3 | *D. p. bogotana* Line SCinv | 135233 | 135464 | 0.20 |
| *D. p. bogotana* Line Suta3 | *D. p. bogotana* Line Toro4 | 134334 | 134335 | 0.00 |
| *D. p. bogotana* Line Suta3 | *D. p. bogotana* Line WhiteER | 135565 | 135880 | 0.37 |
| *D. p. bogotana* Line Suta3 | *D. pseudoobscura* Line S1-A56 | 360524 | 307364 | 4231.23** |
| *D. p. bogotana* Line Suta3 | *D. pseudoobscura* Line S10-A47 | 361629 | 313153 | 3482.49** |
| *D. p. bogotana* Line Suta3 | *D. pseudoobscura* Line S11-A14 | 358637 | 302069 | 4843.21** |
| *D. p. bogotana* Line Suta3 | *D. pseudoobscura* Line S12-M27 | 352897 | 283684 | 7525.26** |
| *D. p. bogotana* Line Suta3 | *D. pseudoobscura* Line S13-A48 | 353294 | 288965 | 6443.23** |
| *D. p. bogotana* Line Suta3 | *D. pseudoobscura* Line S14-A49 | 358728 | 303642 | 4581.23** |
| *D. p. bogotana* Line Suta3 | *D. pseudoobscura* Line S15-A57 | 360959 | 310232 | 3833.82** |
| *D. p. bogotana* Line Suta3 | *D. pseudoobscura* Line S16-A30 | 350890 | 279369 | 8116.11** |
| *D. p. bogotana* Line Suta3 | *D. pseudoobscura* Line S17-M20 | 312747 | 150658 | 56695.21** |
| *D. p. bogotana* Line Suta3 | *D. pseudoobscura* Line S18-M15 | 351123 | 276035 | 8990.09** |
| *D. p. bogotana* Line Suta3 | *D. pseudoobscura* Line S19-A24 | 351877 | 280727 | 8002.36** |
| *D. p. bogotana* Line Suta3 | *D. pseudoobscura* Line S20-M13 | 345586 | 255997 | 13341.78** |
| *D. p. bogotana* Line Suta3 | *D. pseudoobscura* Line S21-M6 | 352231 | 280059 | 8237.99** |
| *D. p. bogotana* Line Toro4 | *D. p. bogotana* Line SCinv | 132939 | 133199 | 0.25 |
| *D. p. bogotana* Line Toro4 | *D. p. bogotana* Line Suta3 | 134335 | 134334 | 0.00 |
| *D. p. bogotana* Line Toro4 | *D. p. bogotana* Line WhiteER | 134961 | 135490 | 1.03 |
| *D. p. bogotana* Line Toro4 | *D. pseudoobscura* Line S1-A56 | 360359 | 307187 | 4235.31** |
| *D. p. bogotana* Line Toro4 | *D. pseudoobscura* Line S10-A47 | 361472 | 313088 | 3470.43** |
| *D. p. bogotana* Line Toro4 | *D. pseudoobscura* Line S11-A14 | 358627 | 302044 | 4846.04** |
| *D. p. bogotana* Line Toro4 | *D. pseudoobscura* Line S12-M27 | 353138 | 283941 | 7515.90** |
| *D. p. bogotana* Line Toro4 | *D. pseudoobscura* Line S13-A48 | 353590 | 289344 | 6419.86** |
| *D. p. bogotana* Line Toro4 | *D. pseudoobscura* Line S14-A49 | 358674 | 303731 | 4557.23** |
| *D. p. bogotana* Line Toro4 | *D. pseudoobscura* Line S15-A57 | 360985 | 310337 | 3821.15** |
| *D. p. bogotana* Line Toro4 | *D. pseudoobscura* Line S16-A30 | 350773 | 279509 | 8057.60** |
| *D. p. bogotana* Line Toro4 | *D. pseudoobscura* Line S17-M20 | 313008 | 151078 | 56501.00** |
| *D. p. bogotana* Line Toro4 | *D. pseudoobscura* Line S18-M15 | 351189 | 276235 | 8954.24** |
| *D. p. bogotana* Line Toro4 | *D. pseudoobscura* Line S19-A24 | 352118 | 280950 | 8000.54** |
| *D. p. bogotana* Line Toro4 | *D. pseudoobscura* Line S20-M13 | 345915 | 256036 | 13420.09** |
| *D. p. bogotana* Line Toro4 | *D. pseudoobscura* Line S21-M6 | 352426 | 280109 | 8267.92** |
| *D. p. bogotana* Line WhiteER | *D. p. bogotana* Line SCinv | 137815 | 137670 | 0.08 |

| | | | | |
|---|---|---|---|---|
| *D. p. bogotana* Line WhiteER | *D. p. bogotana* Line Suta3 | 135880 | 135565 | 0.37 |
| *D. p. bogotana* Line WhiteER | *D. p. bogotana* Line Toro4 | 135490 | 134961 | 1.03 |
| *D. p. bogotana* Line WhiteER | *D. pseudoobscura* Line S1-A56 | 362427 | 307752 | 4460.53** |
| *D. p. bogotana* Line WhiteER | *D. pseudoobscura* Line S10-A47 | 363327 | 313437 | 3677.81** |
| *D. p. bogotana* Line WhiteER | *D. pseudoobscura* Line S11-A14 | 360819 | 302644 | 5101.01** |
| *D. p. bogotana* Line WhiteER | *D. pseudoobscura* Line S12-M27 | 354744 | 284331 | 7758.07** |
| *D. p. bogotana* Line WhiteER | *D. pseudoobscura* Line S13-A48 | 355454 | 289729 | 6695.43** |
| *D. p. bogotana* Line WhiteER | *D. pseudoobscura* Line S14-A49 | 360412 | 304036 | 4783.30** |
| *D. p. bogotana* Line WhiteER | *D. pseudoobscura* Line S15-A57 | 362700 | 310561 | 4037.77** |
| *D. p. bogotana* Line WhiteER | *D. pseudoobscura* Line S16-A30 | 352637 | 279820 | 8383.68** |
| *D. p. bogotana* Line WhiteER | *D. pseudoobscura* Line S17-M20 | 314738 | 151530 | 57127.77** |
| *D. p. bogotana* Line WhiteER | *D. pseudoobscura* Line S18-M15 | 353270 | 276742 | 9295.91** |
| *D. p. bogotana* Line WhiteER | *D. pseudoobscura* Line S19-A24 | 354159 | 281495 | 8306.50** |
| *D. p. bogotana* Line WhiteER | *D. pseudoobscura* Line S20-M13 | 347651 | 256585 | 13724.80** |
| *D. p. bogotana* Line WhiteER | *D. pseudoobscura* Line S21-M6 | 354241 | 280464 | 8575.71** |

** Tajima's relative rate test significant at $p < 0.001$, $X^2$ test, 1 degree of freedom (Tajima 1993).
